# Supporting CIF Format of Biomolecular Structures in GROMACS

**DOI:** 10.1101/2023.09.01.555884

**Authors:** Hengyue Wang, Zhiyong Zhang

**Affiliations:** Department of Physics, University of Science and Technology of China, Hefei, Anhui 230026, China

## Abstract

In the Protein Data Bank (PDB), the PDB format stands as the major file format for protein structures. However, there are certain intrinsic limitations in the PDB format, such as the storage of structural information in a fixed-width format that would be an issue for very large protein complexes. Therefore, the CIF (crystallographic information framework) format has been proposed, which is characterized by superior expansibility. GROMACS, a widely used software suite for molecular dynamics simulations, currently supports only the PDB format. In this study, we modified the source code of GROMACS, which enables it to support the CIF format structure files as input and subsequently generate molecular topology files. This work simplifies the preprocessing of large protein complexes for MD simulations.

**Program summary:** Program title: GROMACS-CIF

GitHub link to program files: https://github.com/zyzhangGroup/Gromacs-CIF

Licensing provisions: GPL-3.0 license

Programming language: C++

Nature of problem: GROMACS currently supports only the PDB format but does not support the CIF format of protein structures. For a very large protein complex, the PDB format is not suitable, and only a CIF file is available in the protein data bank. However, the latter cannot be used as a direct input of GROMACS.

Solution method: Source codes of the program “pdb2gmx” in GROMACS have been modified by adding several CIF analysis functions parallel to the PDB analysis functions.

## Introduction

The Protein Data Bank (PDB) is a widely used repository for protein structure data[1, 2]. It provides two major file formats, namely, the PDB and the CIF (Crystallographic Information Framework) formats.

The PDB format originated in 1971 and is characterized by its strict storage of protein structure data at fixed positions[3, 4]. In a PDB file, the first six characters indicate the type of stored information, such as “HEADER”, “ATOM”, and “HETATM”. The “ATOM” and “HETATM” lines within a PDB file constitute the core content, encompassing details of all heavy atoms within a protein molecule. These data are tightly arranged within a single line. Despite the continued prevalence of the PDB format, it has intrinsic limitations. If the data content exceeds the predetermined limits, the PDB file format becomes invalid. In the beginning, the limit of the PDB format was a trivial issue. However, in recent years, with the development of high-resolution cryo-electron microscopy techniques, an increasing number of structures of large protein complexes have been solved[5]. Such a structure may contain a large number of atomic coordinates that would exceed the limit of the PDB format. Therefore, a new file format is needed to fix this problem.

The CIF format, introduced in 1990[6, 7], has two fundamental distinctions from the PDB format. First, a CIF file is constructed by concatenating data blocks. Second, each data block comprises a label column followed by data columns. The positions of labels in the label column correspond to the positions of data in the data columns, and the data are separated by varying numbers of spaces[4]. The CIF file format offers unlimited expandability, demonstrated by the flexibility to augment the number of data blocks and label columns. Therefore, in 2014, the CIF format replaced the PDB format as the standard format in the PDB[8].

Molecular dynamics (MD) simulation is an important technique for studying the dynamic properties of proteins[9]. Several MD software programs, such as GROMACS[10], AMBER[11], NAMD[12], and CHARMM[13], are available for this purpose. To conduct an MD simulation, an initial protein structure, generally obtained from the PDB, is needed. Taking GROMACS as an example, a preparatory step involves the use of “gmx pdb2gmx” to read atomic coordinates in the PDB file, automatically determine chemical bonds, bond angles, and dihedral angles within the protein molecule, and subsequently generate a GROMACS-formatted coordinate file (gro), a molecular topology file (top), and one or more parameter files (itp) for the following processes.

To the best of our knowledge, until now, GROMACS has lacked support for the CIF format. One solution is to convert the CIF format to the PDB format[14], but this is unsuitable for a large protein complex exceeding the limit of the PDB format. In such a case, one can split the CIF file into multiple PDB files according to the peptide chains, generate separate gro, top, and itp files for each PDB file, and finally merge the files manually. However, this strategy is notably complicated.

As GROMACS is an open-source software, in this work, we have modified its source code. The updated “gmx pdb2gmx” program facilitates direct reading of a CIF file to generate the gro, top, and itp files. This modification simplifies the preprocessing of large protein complexes for MD simulations.

## Program structure

### How GROMACS supports the PDB format

Normally, GROMACS creates topology files from a PDB file through the command “gmx pdb2gmx -f x.pdb -o x.gro -p x.top -i x.itp”. In short, the gro file contains coordinate information in a gmx format; the top file contains information on the molecular topology of the protein; and the itp file contains supplementary information for the topology. Together, they constitute the topology files. Our goal is that GROMACS can run command “gmx pdb2gmx -f x.cif -o x.gro -p x.top -i x.itp” and generate the correct topology files. To enable CIF file support, the source code for how GROMACS supports the PDB file must be found. All related functions and their paths of the source files are listed in Table 1.

**Table 1.**
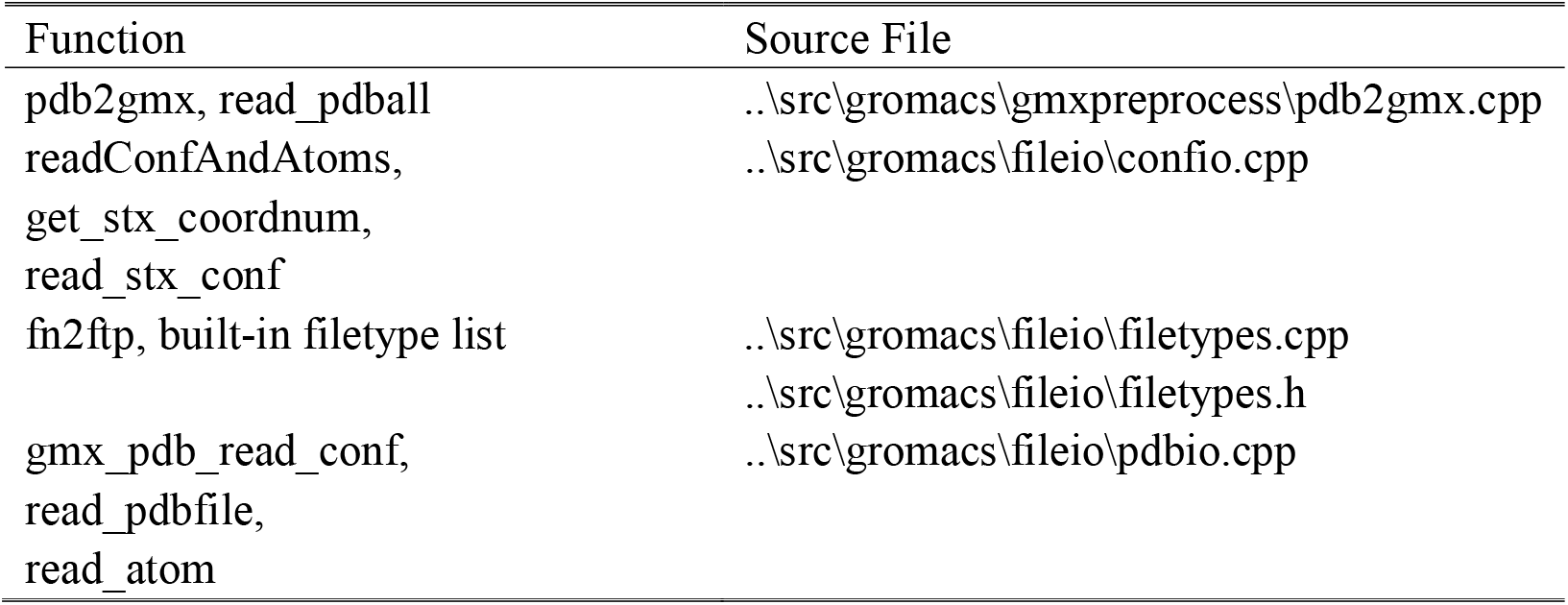
PDB Functions and the Paths of Source Files.

As shown in Figure 1, in the process of the command “gmx pdb2gmx”, the function “pdb2gmx::run” obtains the system path of the PDB file from the user’s input. Then, the path was transmitted through the functions “read_pdball”, “readConfAndAtoms”, “get_stx_coordnum” and “read_stx_conf”.

**Figure 1.**
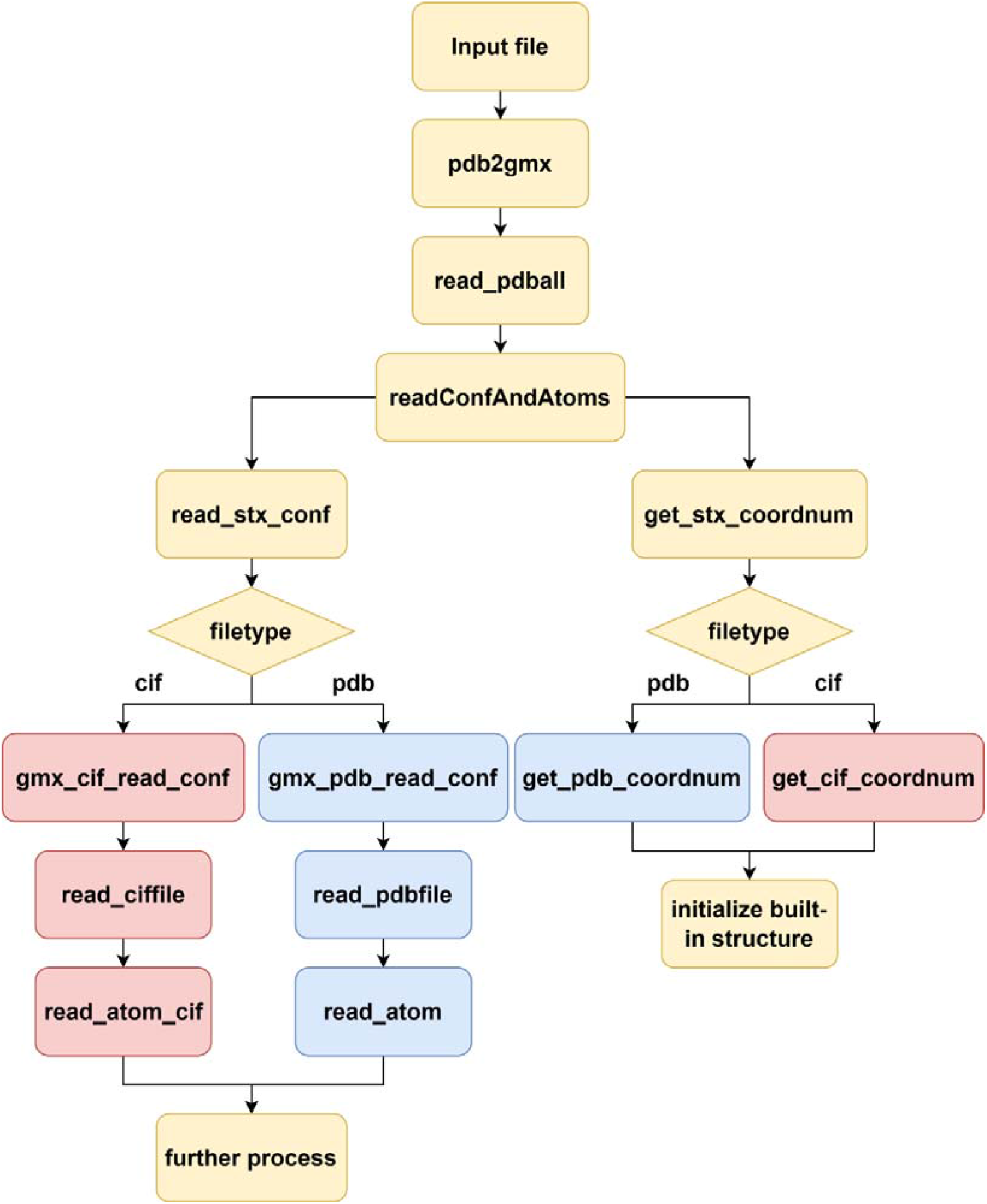
Flowchart of how GROMACS supports the PDB and CIF files. Yellow, the same functions in processing PDB and CIF; Red, the PDB functions; Blue, the CIF functions.

GROMACS not only supports PDB files but also accepts other structure files, such as the gro and g96 formats. To distinguish different file types, GROMACS has a function called “fn2ftp”, which compares the filename extension with the built-in file type list. Therefore, “get_stx_coordnum” and “read_stx_conf” both use “fn2ftp” to identify file types and call corresponding functions. For a PDB file, “get_stx_coordnum” calls the function “get_pdb_coordnum” to acquire the atom number. Then, “get_stx_coordnum” initializes the built-in structure, which will store the whole PDB information. Meanwhile, the PDB file’s system path is transmitted through “read_stx_conf”, “gmx_pdb_read_conf”, and the function “read_pdbfile”. It is worth noting that the PDB file has not yet been analyzed. The functions “read_pdbfile” and “read_atom” will do that.

“read_pdbfile” analyzes the PDB file by the first six characters in each line. For the most important “ATOM” and “HETATM” lines, it calls the function “read_atom” with the whole line as input for further processing. In “read_atom”, the whole line is strictly split by bits due to the rule of PDB file (Table 2). Then, each data part is stored in the corresponding variable contained by the built-in structure initialized in “get_stx_coordnum”.

**Table 2.** Corresponding relationship between the PDB and CIF format. The same data are stored both at the fixed location in the PDB file and label in the CIF file.

| Data Meaning | Example | Location in PDB | Label in CIF |
| --- | --- | --- | --- |
| Line type | ATOM | 1~6 | _atom_site.group_PDB |
| Atom number | 986 | 7~11 | _atom_site.id |
| Atom name | CD1 | 13~16 | _atom_site.label_atom_id |
| Residue name | TRP | 18~21 | _atom_site.label_comp_id |
| Chain ID | A | 23 | _atom_site.label_asym_id |
| Residue number in chain | 126 | 25~28 | _atom_site.auth_seq_id |
| X location | 31.655 | 33~40 | _atom_site.Cartn_x |
| Y location | -2.030 | 41~48 | _atom_site.Cartn_y |
| Z location | -6.788 | 49~56 | _atom_site.Cartn_z |
| Weight | 1.00 | 57~62 | _atom_site.occupancy |
| B-factor | 23.24 | 63~69 | _atom_site.B_iso_or_equiv |
| Element name | C | 80~81 | _atom_site.type_symbol |

### Comparison between the PDB and CIF format

Table 2 illustrates the corresponding relationship between the PDB and the CIF file through an example (PDB ID 2LZM[15]). Obviously, all information in the PDB file is also contained in the CIF file. Our idea is that if we store the corresponding CIF file’s data into those variables in the function “read_atom”, GROMACS will recognize them as the PDB input and continue its work. Therefore, we could create a topology file directly from the CIF file by modifying the data-reading process in GROMACS.

### How to support the CIF format in GROMACS

First, we added the CIF file type support in the source files “filetypes.cpp” and “filetypes.h”. Then, we modified “get_stx_coordnum” and “read_stx_conf” to correctly identify the CIF file type and call the corresponding CIF functions. Next, we added the CIF functions “get_cif_coordnum”, “gmx_cif_read_conf”, “read_ciffile” and “read_atoms_cif” to the source file “pdbio.cpp”. These new functions are parallel with the PDB functions, as shown in Figure 1 and Table 3.

**Table 3.** Usage of the PDB and CIF functions.

| PDB Function | CIF Function | Usage |
| --- | --- | --- |
| get_pdb_coordnum | get_cif_coordnum | Get number of all atoms |
| gmx_pdb_read_conf | gmx_cif_read_conf | Open structure file |
| read_pdbfile | read_ciffile | Analyze structure file, and separate out lines containing atom information for further usage |
| read_atom | read_atom_cif | Analyze atom information line, and store useful part in corresponding variable |

A CIF file is divided into several information boxes by “#”. We only need the box containing PDB’s “ATOM” and “HETATM” line information, which could be easily located by its unique label “_atom_site.group_PDB”. To analyze the CIF file in the function “read_ciffile”, we first search the whole CIF file to find the unique label. Then, we check all labels and save only the locations of labels shown in Table 2. Last, in the function “read_atoms_cif”, we divided the line by whitespace. Because the location of the data and label is the same, we could store the data by its location in exactly the same variable as in the function “read_atom”. For more details, check the source codes in the Supplementary Material.

### Program usage

To verify the correctness of gro, top and itp files generated directly from the CIF file, we calculate their md5 values and compare them with topology files generated from the PDB file.

For a simple case, we selected a relatively small protein, the bacteriophage T4 lysozyme (PDB ID 2LZM[15]). It is a single-chain protein with only 164 amino acid residues and 1309 heavy atoms (Figure 2a). The structure database provides both the PDB and the CIF files of the T4 lysozyme. Two groups of topology files are generated by the modified GROMACS using the PDB and the CIF file as input. Then, we calculate the md5 value of the data section in the topology files. The results from the PDB and the CIF file are identical, so we prove the correctness of the CIF-generated topology files for small proteins.

**Figure 2.**
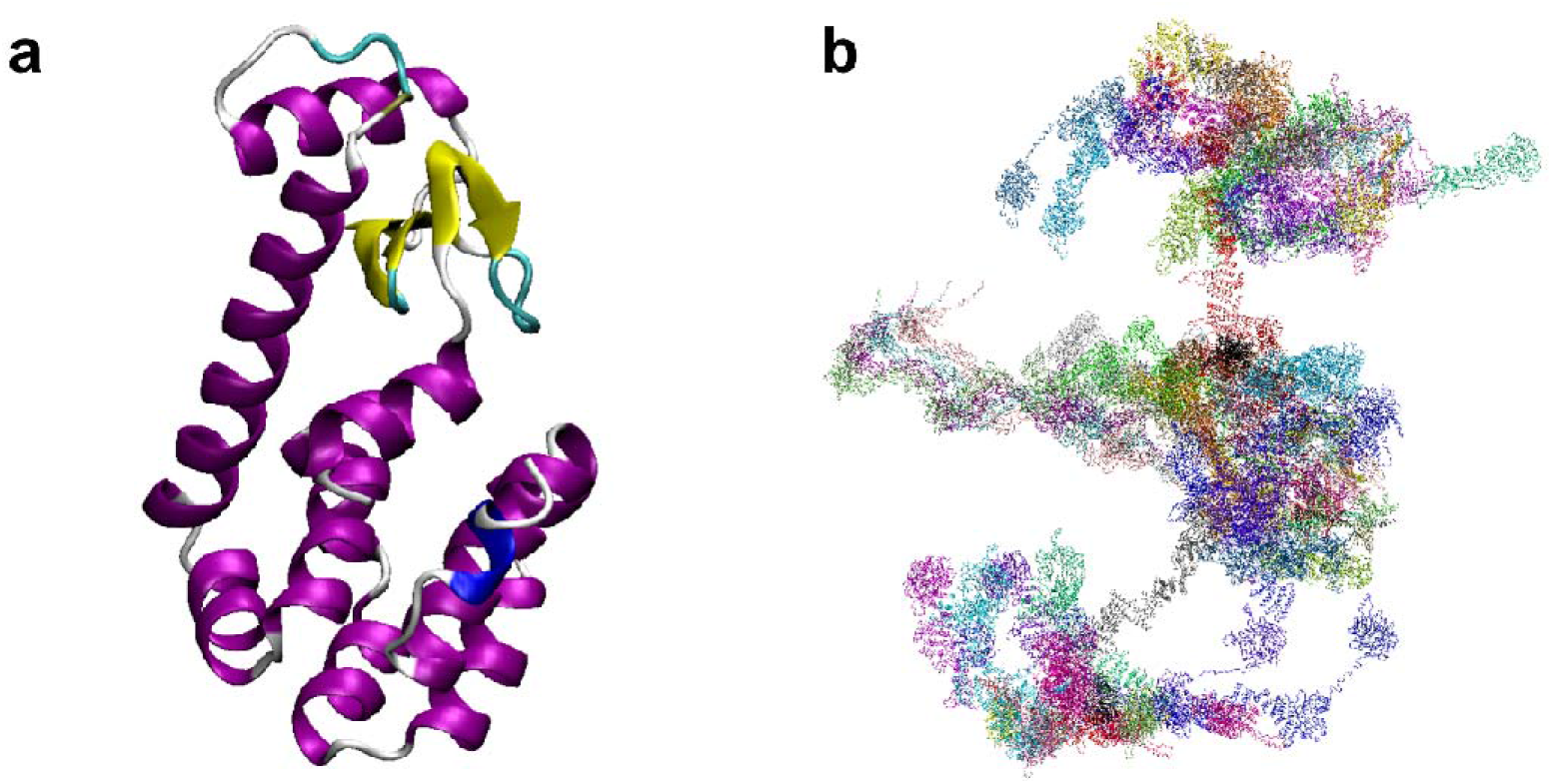
The systems used to test the modified GROMACS. (a) The bacteriophage T4 lysozyme (2LZM). (b) The dilated human nuclear pore complex (7R5J). Since the whole complex has C8 symmetry, only one-eighth of the structure is included in the CIF file.

For a complicated case, we chose the dilated state of the human nuclear pore complex (PDB ID 7R5J[16]). It contains 25 different nucleoporins and 101 peptide chains with 617133 heavy atoms in total (Figure 2b). This large protein complex obviously exceeds the limit of the PDB format, so that the structure database only provides a CIF file and PDB format-like files, and the latter are several PDB files, each including parts of the data to circumvent limitations.

The modified GROMACS can successfully generate topology files through 7R5J.cif. Next, we selected 5 different peptide chains in 7R5J. The same data in PDB format-like files are merged into one PDB file as the input of the modified GROMACS to generate topology files. As in the case of 2LZM, we separated the two groups of the topology files’ data section out and proved identical through the md5 values.

In conclusion, the modified GROMACS could generate correct topology files from the CIF files.

## Conclusion

In this work, we modified the open-source software GROMACS to enable support for the CIF file format, allowing for the direct utilization of CIF files as input to generate topology files of proteins. To validate the accuracy of the generated topology files, we selected two systems: a relatively small protein (bacteriophage T4 lysozyme) and a large protein complex (asymmetric unit of the dilated human nuclear pore complex). A comparison between topology files generated from the CIF format and those generated from the PDB format was conducted, thereby affirming their correctness. For very large protein complexes lacking PDB-format files, our contribution streamlines the preprocessing phase of molecular dynamics simulations.

## Acknowledgment

This work is supported by the National Key Research and Development Program of China (2021YFA1301504), the National Natural Science Foundation of China (91953101), and the Chinese Academy of Sciences Strategic Priority Research Program (XDB37040202). We are grateful to Mr. Yundong Zhang for technical support and to Zhiqiang Wang, Chaomei Yan, and Shuming Wang for helpful discussion.

## Supplementary Material

Our installation guidance, source codes and examples are available at https://github.com/zyzhangGroup/Gromacs-CIF.

